# Causal affordances shape Hornbills’ problem-solving strategies

**DOI:** 10.1101/2025.04.03.646957

**Authors:** Elias Garcia-Pelegrin

## Abstract

Insight—that eureka moment when understanding emerges without trial-and-error—has traditionally been limited to great apes, corvids, and parrots (Byrne, 2016; Emery & Clayton, 2004; Lambert et al., 2019). The vertical string-pulling paradigm, a hallmark test of avian cognition (Jacobs & Osvath, 2015), requires birds to retrieve suspended food by incrementally pulling the string with their bill while securing it with their foot—demonstrating both motor coordination and causal understanding. However, Oriental pied hornbills (*Anthracoceros albirostris*) present an interesting case due to their unique anatomical constraints—their fused proximal phalanges prevent them from employing the typical foot-assisted string-pulling technique used by other birds. This study challenged six Oriental pied hornbills with two distinct vertical string-pulling problems, each requiring different solving strategies based on their causal properties. Despite their pedal limitations, five of the six subjects successfully solved both problems with remarkable efficiency, achieving solutions within an average of 17.6 seconds of their first encounter with the tasks. Most notably, the successful hornbills immediately employed task-specific biomechanical strategies that directly addressed the causal relationship between the string and the food reward: vigorous shaking movements to dislodge unsecured rewards, versus precise bill-and-tongue manipulation for rewards firmly attached to the string. The hornbills’ immediate application of causally appropriate, task-specific solutions without prior trial-and-error strongly supports the genuine insight hypothesis rather than associative learning explanations. These findings reveal sophisticated problem-solving capabilities in this previously understudied taxon and contribute valuable evidence to our understanding of convergent cognitive evolution across phylogenetically distant species.

## Main Text

The concept of “insight” has remained contentious since Köhler’s foundational work with chimpanzees (Pan troglodytes)(Köhler, 1925). While Köhler documented apparent spontaneous understanding when subjects stacked boxes to reach food or combined sticks to create tools, subsequent critical analyses have challenged these interpretations. Researchers noted the chimpanzees’ initial unsuccessful attempts and proposed that prior experience with task components likely facilitated their problem-solving (Birch, 1945; Schiller, 1952; Shettleworth, 2012). Analogous findings with pigeons (Epstein et al., 1984) provide further evidence that purported demonstrations of insight may instead represent cumulative associative learning rather than genuine cognitive breakthroughs characterized by sudden understanding.

The act of pulling a vertically suspended string demands of the subject to partake in some degree of motor planning effort, as any untimed release of the string would result in the loss of the reward. Corvids and parrots typically solve this problem within seconds of exposure to it (Heinrich & Bugnyar, 2005; Lambert et al., 2019; Werdenich & Huber, 2006), using a complex technique of reaching, gripping, pulling, looping, and stepping on the string repeatedly until obtaining the reward. Within this context, an ‘insight’ hypothesis would suggest that birds mentally visualize how these actions affect the food’s position (Shettleworth, 2012). However, debate also exists about whether this demonstrates true causal understanding. Some argue the behaviour is merely reinforced by seeing the reward move closer with each pull (Taylor et al., 2012). Indeed, when visual feedback is removed, such as by hiding the string’s end in an opaque tube, New Caledonian Crows *(Corvus moneduloides)* fail to produce the characteristic motor operation (Taylor et al., 2010).

Stepping on the string to prevent slippage is a critical challenge in string pulling. This action may relate to foraging habits where birds secure items underfoot to free their beaks (Seibt & Wickler, 2006). However, the correlation is inconsistent - some birds that use feet while feeding fail at string pulling, while others that don’t use feet during feeding succeed at the task (see (Jacobs & Osvath, 2015) for a review). Arguably more important than how the animal uses its feet is the actual morphological nuances of their appendages that allow for the use of the feet in such a manner. Birds possess varied foot structures adapted to different functions. Corvids with anisodactyl feet and parrots with zygodactyl feet allow for a pincer-like grip (Abourachid et al., 2017), enabling them to tightly step on a string after pulling it to prevent it from slipping back. This is crucial as not all avian pedal configurations allow for this specific and integral requirement for the string-pulling manoeuvrer. Hornbills (Bucerotidae, order Bucerotiformes) display extreme syndactyly, which results in the fusion of the proximal phalanges of their three front toes (Kemp, 1995). This configuration restricts their ability to generate the necessary biomechanical force to effectively step on a string. Therefore, hornbills serve as an interesting and apt model for investigating how physical limitations can drive the development of alternative cognitive strategies in tool use and problem-solving.

Both vertical string-pulling problems presented to the hornbills could be solved using the “stepping on the string” manoeuvre typically exhibited by corvids and parrots. However, the tested hornbills quickly solved both problems despite lacking this capability. One of the problems, the unsecured food problem (Figure 1, *left*), consisted of an 8 cm^3^ circular container with a looped handle connected to a string hanged from a perch. The container was open at the top, and inside it the food rewards where left loose. Instead, the secured food problem (Figure 1, *right*), consisted of a single pellet (1 cm) tightly secured to the end of a string using vinyl electrical tape. The two problems were tested one after the other, with a 5-minute break in between, and a counterbalanced order to mitigate any learning effects. Each hornbill was given one attempt per problem, with up to 15 minutes available to solve each.

**Figure 1.**
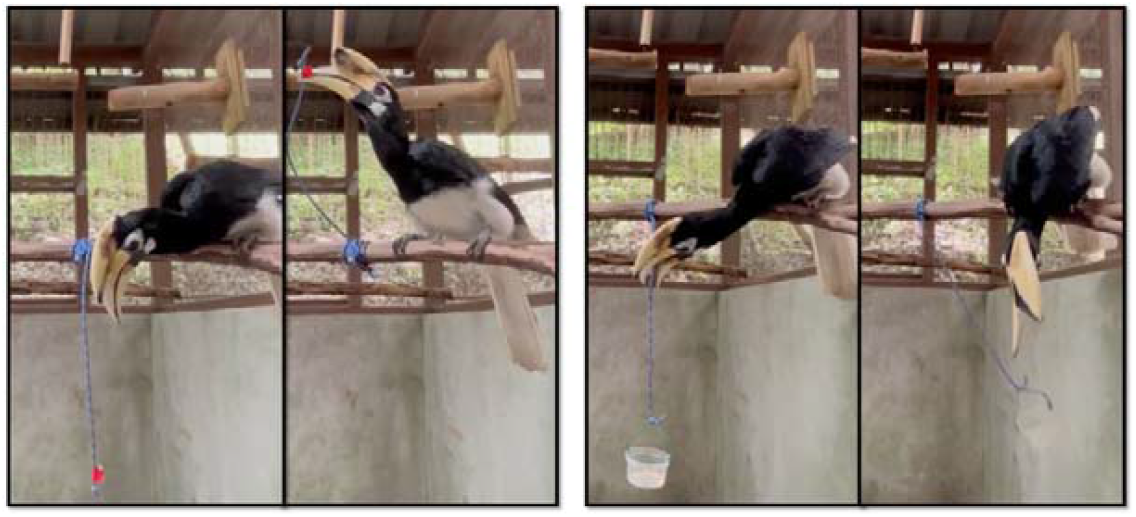
Experimental setup for vertical string-pulling tasks (left) secured food condition and (right) unsecured food condition.

## Results and Discussion

All hornbills approached both problems immediately after starting the trial (mean latency 1.03 seconds for secured condition, and 0.46 seconds for unsecured condition). Whilst neither of the hornbills tested had ever encountered a set up like this one before, the low latency to approach is unsurprising given that Asian hornbills have typically low levels of neophobia and high levels of food motivation (Garcia□ Pelegrin et al., 2022). Latency to solve both problems was relatively fast, with no significant difference between them (mean latency 18.1 seconds for secured condition, and 17.8 seconds for unsecured condition). Notably, the speed at which these string-pulling tasks were solved is comparable to the speeds usually documented for naïve corvids and parrots that solve on the first trial (e.g., 9 – 15 s for keas *(Nestor notabilis)* (Werdenich & Huber, 2006) and 6–37s for New Caledonian crows (Taylor et al., 2012)).

The analysis of the hornbills’ solving techniques used identified two behavioural strategies for problem-solving. These strategies varied based on the problem type; one was used by all subjects to address the unsecured string problem, while the other was adopted by 5 out of 6 subjects for the secured condition. In the unsecured condition, the hornbills repeatedly grasped and released the string, shaking it vigorously to dislodge the food from the container (Video S1). They would then go to the floor where the food had landed, eat it, and repeat this process until all the food was dislodged. For the secured condition, all but one subject immediately took a different approach, using their bills and tongues to manoeuvre the entire string, pulling the food closer and then dislodging it by applying sideways tension (Video S2). One hornbill attempted to solve the secured condition using the same shaking method from the unsecured task, and consequently never succeeded.

Subjects exhibited goal-directed behaviour, favouring efficient over exploratory actions (Figure 2). Rewards were obtained exclusively through shaking motions for unsecured conditions and direct pulling for secured conditions. Subjects manipulated the apparatus using only their bills, never their feet. Efficiency ratios (efficient actions/total actions) remained consistently high across conditions: secured condition (M=0.82, SD=0.08), and unsecured condition (M=0.84, SD=0.09) (Figure 2). The hornbills displayed consistent behavioural repertoires across individuals, with only minor technique variations. Those encountering the secured condition second immediately shifted from shaking to pulling, while two of three hornbills that started with the secured condition directly employed shaking without first attempting to pull.

**Figure 2:**
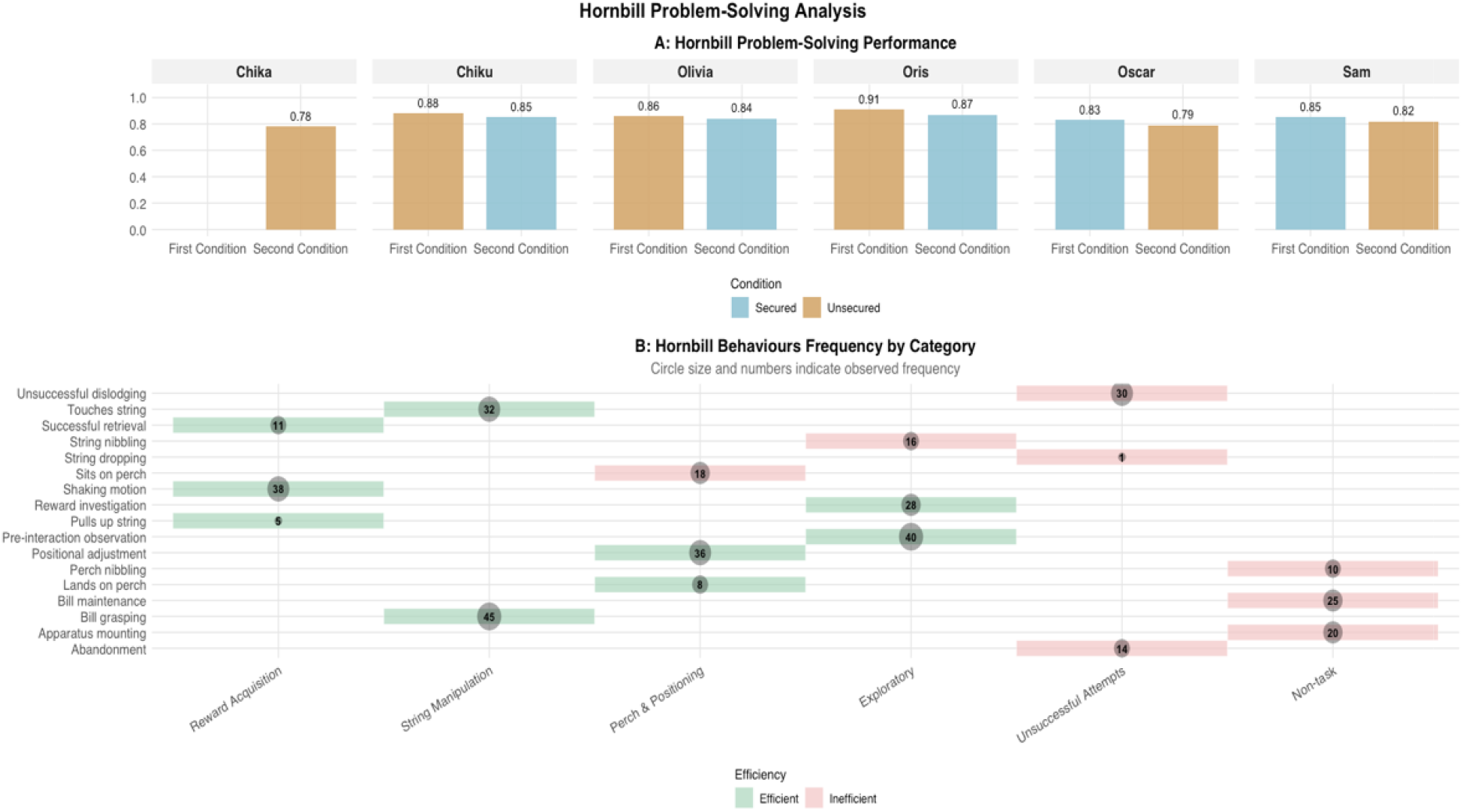
Hornbill problem-solving performance and behavioural frequencies across experimental conditions. ***Graph A:*** Efficiency ratios (efficient actions / total actions) of subjects under secure and unsecure conditions. ***Graph B***: Frequency of observed behaviours categorized as efficient (green) or inefficient (red) across behavioural categories. Black dots represent the overall frequency of behaviours within each category.

The fact that they were naïve to both problems and quick in applying the behaviour necessary to succeed limits any explanation about trial-and-error learning. Interestingly, the speed at which they solved these problems is comparable to that reported for corvids and parrots, even though these use the stepping motor pattern. Evidently, the stepping manoeuvre would be more advantageous for the unsecured food condition than the hornbill’s technique, as it would allow access to the contents of the cup in one go, rather than requiring multiple shakes to dislodge all the food. In this study, a problem was considered solved once the first food was consumed, not when all the food was accessed. Conversely, the hornbill’s method for the secured condition proves to be more efficient than the stepping manoeuvre, reaching the food with minimal motor effort and significantly faster. However, given that hornbills possess a serrated bill, this action is likely species-specific and not generalizable to most other bird species.

The problem-solving techniques exhibited by the hornbills seemed to capitalize on the unique affordances provided by the relationship between the string and how the food was attached to it. Crucially, while both problems could have been solved using the stepping behaviour, employing the shaking method used for unsecured food would not successfully grant access to the reward in the secured food condition, as it was firmly attached to the string. Similarly, without appendages to free the bill for eating, using the combination of bill and tongue manoeuvres to access the reward used in the secured condition for the unsecured scenario would leave the animal with the cup in its bill but unable to consume the food it contained.

To challenge the insight hypothesis, it is often suggested that motor patterns employed in string tasks might be reinforced by association. When an agent pulls the string and sees the reward move closer, this visual cue strengthens the behaviour until the reward is acquired (Taylor et al., 2010). However, in the unsecured condition, the shaking the string technique employed makes the reward move away from the subject. Moreover, if the shaking action was reinforced through trial and error, hornbills successful in this, which also received this condition first, should briefly repeat it in the secured condition before switching techniques after failure. Yet, all three hornbills who started with the unsecured condition succeeded in the secured condition immediately, without ever letting go of the string. Similarly, none of the hornbills that started with the secured condition, brought the reward closer to themselves at any point in their unsecured condition.

Consequently, considering the speed of resolution, the lack of trial-and-error learning, and the fact that the subjects utilized specific techniques tailored to each problem, this report provides convincing evidence that this relatively understudied bird species can account for the causal variables of physical problems and plan their behaviour accordingly before interacting with them in a manner typically described as insight. Recent findings indicate that Oriental pied hornbills also exhibit advanced levels of object permanence, a cognitive complexity typically attributed exclusively to corvids, parrots, and great apes (Yao & Garcia-Pelegrin, 2024). In conclusion, the patterns documented in this report demonstrate that this species possesses sophisticated cognitive abilities previously unrecognized in the scientific literature. These findings not only expand our understanding of avian intelligence but also suggest the need to reassess how we evaluate and classify cognitive capabilities across the avian taxa. Further research may reveal additional examples of advanced cognition in this understudied avian species.

## Materials and Methods

### Subjects

Subjects were 6 oriental-pied hornbills (*Anthracoceros albirostris*), comprising three females and three males. All subjects were housed at the Mandai Wildlife Reserve Collection, either individually (if unpaired) or in pairs within aviaries measuring 6m × 4m × 3m (width × length × height). The birds maintained their regular feeding schedule throughout the study, receiving a standard diet of fresh fruits and nutrient pellets with continuous access to water. Prior to this study, none of the subjects had experience with string-pulling tasks. One experimenter (EG-P) collected all the data.

### Apparatus

The experimental setup consisted of a standardized apparatus featuring a 1-meter string attached to a perch via a carabiner. This configuration prevented subjects from accessing rewards from the enclosure floor. Two variants of the apparatus were used, differentiated by the method of reward attachment.

#### Vertical problem 1. Unsecured food

In this problem, the end of the string was connected to an 8 cm^3^ circular container with a looped handle. The container was open at the top, and inside it were either 10 hornbill food pellets, each about 1 cm long, or 5 papaya pieces measuring 2 cm each.

#### Vertical problem 2. Secured food

In this problem, a single pellet (1 cm) was firmly attached to the end of a string using vinyl electrical tape. Since it was crucial that the pellet remained securely fixed to the string (unlike in the first problem), several vigorous shake tests were manually conducted by the experimenter before each testing round.

### Procedure

Each subject underwent testing for both conditions in a single session, with conditions presented in a pseudorandomized, counterbalanced order. For each trial, the experimenter first set up the apparatus in the testing chamber while the subject remained in a separate area. The trial began when the hornbill was permitted to enter the testing chamber, ensuring standardized starting conditions across all trials. The trial began when the hornbill was permitted to enter the testing chamber, ensuring standardized starting conditions across all trials. A 5-minute break separated the conditions, with each trial lasting up to 15 minutes.

### Analysis

All sessions were video-recorded and analysed using BORIS software version 8.22.16 (Friard & Gamba, 2016). The analysis focused on initial approach latency, problem-solving strategies, success rates, and time to solution. Five of six subjects successfully solved both conditions on their first attempt, while one male subject (Chika) solved only the unsecured food condition.

#### Latency to approach and solve

Latencies were coded from video recordings using a systematic approach with clearly defined start and end points. For approach latency, timing began at the moment the experimental apparatus was fully set up and the experimenter stepped away from the testing area. Timing ended when the subject made first physical contact with any part of the apparatus.

Latency to solve was measured as the duration between the same starting point as latency to approach and successful retrieval of the food reward. For unsuccessful attempts (as in Chika’s case with the secured condition), the session was terminated after 15 minutes of engagement without success, and N/A was recorded.

All latency measurements were coded independently by two observers who were blind to the study hypotheses to prevent observer bias. Inter-observer reliability was calculated using intraclass correlation coefficients (ICC), yielding high agreement for both approach latency (ICC = 0.97, p < 0.001) and solution latency (ICC = 0.99, p < 0.001). Any discrepancies greater than two seconds between observers were resolved through frame-by-frame video analysis and discussion to reach consensus.

## Ethics

The experimental protocol received approval from both the National University of Singapore’s Institutional Animal Care and Use Committee (IACUC protocol number R23-0737) and the Mandai Wildlife Reserve research panel.

## Acknowledgments

I sincerely thank Mandai Wildlife Group for providing access to their facilities and for their invaluable assistance throughout my research. I am especially grateful to the dedicated team at Animal Behaviour and Programmes for their essential assistance in this study. A special thanks to Gail Laule and Kelly Chew for their support which has been instrumental in completing this work.

